# Enhancing Fairness in Disease Prediction by Optimizing Multiple Domain Adversarial Networks

**DOI:** 10.1101/2023.08.04.551906

**Authors:** Bin Li, Xinghua Shi, Hongchang Gao, Xiaoqian Jiang, Kai Zhang, Arif O Harmanci, Bradley Malin

## Abstract

Predictive models in biomedicine need to ensure equitable and reliable outcomes for the populations they are applied to. Unfortunately, biases in medical predictions can lead to unfair treatment and widening disparities, underscoring the need for effective techniques to address these issues. To enhance fairness, we introduce a framework based on a Multiple Domain Adversarial Neural Network (MDANN), which incorporates multiple adversarial components. In an MDANN, an adversarial module is applied to learn a fair pattern by negative gradients back-propagating across multiple sensitive features (i.e., characteristics of individuals that should not be used to discriminate unfairly between individuals when making predictions or decisions.) We leverage loss functions based on the Area Under the Receiver Operating Characteristic Curve (AUC) to address the class imbalance, promoting equitable classification performance for minority groups (e.g., a subset of the population that is underrepresented or disadvantaged.) Moreover, we utilize pre-trained convolutional autoencoders (CAEs) to extract deep representations of data, aiming to enhance prediction accuracy and fairness. Combining these mechanisms, we alleviate biases and disparities to provide reliable and equitable disease prediction. We empirically demonstrate that the MDANN approach leads to better accuracy and fairness in predicting disease progression using brain imaging data for Alzheimer’s Disease and Autism populations than state-of-the-art techniques.

## 1. Introduction

Precision medicine represents a tailored approach to healthcare, where treatments and interventions are customized to the individual’s unique genetic, environmental, and lifestyle factors. This personalized approach has the potential to enhance both the effectiveness and fairness of medical care. By considering the specific characteristics of each individual, precision medicine can help to mitigate biases that may arise from a “one-size-fits-all” approach, where certain populations might be underrepresented or disadvantaged.

The utilization of artificial intelligence (AI), particularly deep learning (DL), in the analysis of biomedical data, has emerged as a promising approach for improving healthcare outcomes. However, there are growing concerns that the blind application of such methods can induce harm and perpetuate inequities.^1^ Biases present in both data and AI models can reinforce disparities in healthcare applications, leading to unequal outcomes across different patient subpopulations. However, it is crucial to distinguish between bias and fairness. Bias refers to the systematic error or deviation in predictions or estimates as a result of the model’s assumptions. Fairness, on the other hand, relates to the equitable treatment of all individuals, regardless of their membership in certain subpopulations. However, mitigating biases could reduce disparities, making predictions more equal across different groups. Therefore, ensuring fairness in AI involves creating models that provide accurate diagnostics and unbiased treatment decisions, irrespective of the patient’s subpopulation, thereby promoting equitable access to healthcare.^2^

However, it is difficult to mitigate biases to achieve fair AI/DL for biomedical data analysis.^3^ One of the key difficulties is identifying and quantifying biases within the data. Once these biases are identified, an additional challenge arises: preventing these biases from exacerbating disparities in subsequent AI/DL tasks. Biomedical datasets are often prone to biases that arise from imbalances in the demographic composition, socioeconomic factors, and disparities in healthcare practices across subpopulations.^4^ As a result, predictions based on AI/DL that are optimized on such data often lead to misleading or wrong findings that can endanger minority groups.

Various techniques have been proposed to address fairness concerns for AI/ML in biomedicine. Approaches to bias mitigation typically fall into three gross categories associated with the pre-processing, in-processing, and post-processing of an ML model.^5^ In preprocessing, inequities in data are removed prior to model training.^6^.^7^ During in-processing, the model training process is performed to actively mitigate discrimination during model training.^8,9,10^ In post-processing, the output of a trained model is adjusted to achieve fairness.^11,12^ Pre-processing can be performed by resampling existing data, incorporating new data, or adjusting data labels. In-processing methods use adversarial techniques, impose constraints and regularization, or ensure fairness of underlying representations during training. Finally, post-processing entails group-specific modification of decision thresholds or outcomes to ensure fairness in the application of model predictions. Different approaches may be optimal depending on the setting and stage of model development.^13,14^

In this paper, We introduce the notion of a Multiple Domain Adversarial Neural Network (MDANN), which is designed to enhance fairness by mitigating biases from multiple sensitive features simultaneously. Note that sensitive features in this context refer to characteristics of individuals (e.g., race, gender, age, etc.) that should not be used to discriminate unfairly between individuals when making predictions or decisions. There are several notable contributions to bias mitigation and fairness enhancement in biomedical data analysis. **First**, we extract deep representations of input data from the embedding layer of a pre-trained convolutional autoencoder (CAE). These deep representations contain enriched information within higher-dimensional spaces. This, in turn, significantly improves prediction accuracy and fairness in disease prediction tasks. Additionally, we show that the CAE is a powerful feature extractor, contributing to enhanced prediction fairness. **Second**, we employ an AUC-induced minimax loss function that takes into account the Area Under the Receiver Operating Characteristic Curve (AUC) score, in contrast to the conventional accuracy-induced loss, such as cross-entropy. As we empirically illustrate, the minimax loss function outperformed the standard cross-entropy loss in handling imbalanced data, leading to improved classification performance and fairness in predictions for the minority class. **Finally**, we investigated the impact of adversarial modules on prediction performance in the context of bias mitigation using the MDANN. By addressing multiple sensitive features, we demonstrate that the introduction of adversarial components effectively enhances fairness in disease prediction.

## 2. Related Work

### 2.1. Fairness improvement in machine learning

Fairness enhancement and bias mitigation in DL has been addressed through various methodologies. Pre-processing techniques that re-weight or re-sample instances to balance representation across groups have been employed, but these may overlook latent or structural biases.^15^ Algorithmic fairness interventions, such as incorporating fairness constraints or regularization terms during model training, provide nuanced control over fairness but may compromise model accuracy or generalization^16^). Recent advancements include the debiasing of embeddings^17^ and the certification of disparate impact.^18^ These are sophisticated solutions, but often require complex optimization. Investigations into the long-term impact of fairness interventions^19^ and the trade-offs between fairness and accuracy^16^ have further enriched the understanding of this complex field. Comprehensive surveys have further elucidated these methods and their multifaceted implications^20^.^21^

### 2.2. Domain adversarial neural network

Domain Adversarial Neural Networks (DANN) have demonstrated significant advancements in domain adaptation, particularly in scenarios where models are trained on a source domain and applied to a disparate target domain.^22^ Essentially, a domain Adversarial Network (DANN)^22^ is a machine learning technique used for domain adaptation, a scenario where a model is trained on a source domain and then applied to a different target domain. DANN was originally proposed as a way to address the problem of domain shift, where the distributions of the source and target domains differ significantly, leading to a drop in model performance when applied to the target domain.^23^ The primary goal of a DANN is to learn domain-invariant representations between the source and target domains. This is achieved by adding a domain classifier to the model, which is tasked with predicting the domain of the input data (i.e., source or target). The domain classifier is trained to make accurate domain predictions, while the main task of the model (e.g., classification or regression) is trained to make accurate predictions on the source domain data.^24^

Transitioning from the theoretical underpinnings of DANN to its practical applications, it’s important to note that biomedical datasets often exhibit significant variation across different sources, institutions, modalities, and populations. In this context, DANN’s ability to handle domain adaptation becomes particularly valuable. It allows a model trained on one domain (source) to be applied to a different domain (target) without significant loss in performance. This is especially beneficial in biomedical contexts where data may be collected from various hospitals, instruments, or demographic groups.^25^ While a DANN can address domain shift, it may not improve fairness related to patient demographics, medical conditions, or data collection protocols.^26^ Furthermore, ethical considerations and the need for careful validation in biomedical applications add layers of complexity that may constrain the effectiveness of a DANN in fully addressing fairness. These limitations underscore the need for continued research and innovation in adapting DANNs for the nuanced requirements of fairness in biomedicine. In the context of fairness enhancement, a DANN can be expanded to multiple-domain adversarial networks, as demonstrated in this study, to address fairness concerns when working with multiple sensitive domains. This is particularly relevant in real-world applications where machine learning models might be applied to data gathered from various groups and fairness needs to be maximized for all domains and groups.

## 3. Methods

In this section, we introduce the MDANN approach for fairness enhancement in disease prediction. Source codes are available at https://github.com/shilab/MDANN. **Fig. 1** depicts the architecture of the MDANN. The MDANN architecture is tailored to the unique demand for fairness enhancement and is composed of three fundamental and synergistic modules (shown in bold): **1) Feature Extractor:** A pre-trained convolutional autoencoder designed to delve into the complex structure of biomedical data, extracting deep and meaningful representations that serve as the foundation for subsequent calculation. **2) Adversarial Module:** A novel and innovative approach to improve fairness, utilizing multiple adversarial networks to learn fair patterns across various domains, reflecting our commitment to equitable treatment across diverse demographic groups. **3) AUC-induced Min-max Loss Function:** A carefully designed loss function that focuses on the AUC, providing a robust and sensible metric for imbalanced data classification, and underscoring our nuanced understanding of the challenges posed by imbalanced biomedical data. In summary, these three modules form an integrated and end-to-end learning network, each contributing unique strengths and working in harmony to extract informative representations, mitigate multiple biases, and promote fairness against label imbalance problems. In the following sections, we will introduce each module, detailing their design and functionality within the MDANN framework.

**Fig. 1.**
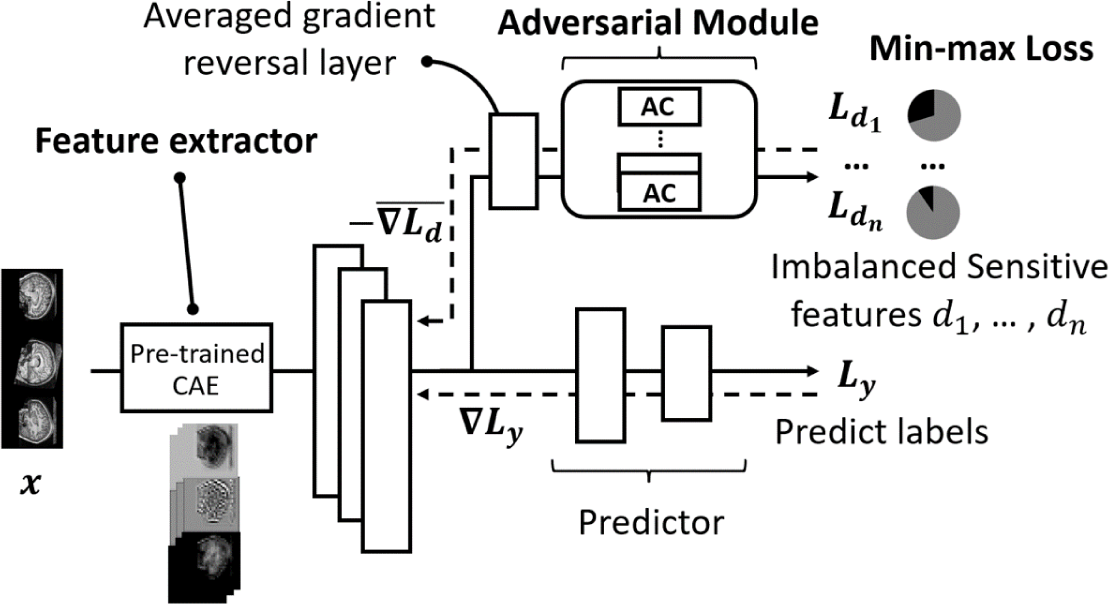
The MDANN architecture. Three fundamental modules have been incorporated into the MDANN including a feature extractor, an adversarial module and an optimizer. The feature extractor is essentially a pre-trained convolutional autoencoder (CAE) to extract the embedding content as a deep representation from imaging samples. The adversarial module contains several adversarial components (ACs) which are used to learn distributions of multiple sensitive features simultaneously and pass back the negative gradients through an averaged gradient reversal layer. The optimizer is based on the min-max function with an AUC-induced minimax loss to address the label-imbalance problem to mitigate biases in input data.

### 3.1. Feature Extractor

The feature extraction process employed in this study entails the utilization of a pre-trained convolutional autoencoder (CAE) model, consisting of convolutional layers serving as the encoder and deconvolutional layers acting as the decoder. The CAE model is designed to acquire underlying patterns in the data to generate images resembling the original input images. Within the context of the MDANN framework, deep representations of imaging samples are extracted from the last convolutional layer (bottleneck) of the model. This approach facilitates the transformation of 2D images into high-dimensional vectors in latent space through multiple convolutional calculations. The resulting deep representations possess an enriched level of grayscale and texture details, rendering them conducive to mitigating the influence of biased information. By adopting this feature extraction methodology, our research endeavors to enhance the fidelity and informativeness of the extracted representations, ultimately contributing to the reduction of biases and promoting fairness in the subsequent stages of predictive modeling.

### 3.2. Adversarial module

The adversarial module within our framework is implemented through the incorporation of adversarial components, each composed of two linear layers acting as predictors. These components are seamlessly integrated with the feature extractor by employing a gradient reversal layer, which exerts its influence during backpropagation-based training. Specifically, the gradient reversal layer operates by multiplying the averaged negative gradient, thereby facilitating the adversarial alignment of feature distributions across biased groups. The primary objective of the adversarial module is to achieve robust adversarial learning, wherein the feature distributions pertaining to different sensitive attributes are rendered as indistinguishable as possible for each component.

A significant contribution of our approach lies in the simultaneous mitigation of multiple biases through the use of multiple adversarial networks. This innovation enables the MDANN to effectively address biases across various sensitive features simultaneously, as opposed to traditional approaches that may tackle biases in a sequential or isolated manner. By incorporating multiple adversarial components, each catering to a distinct sensitive attribute, the model is endowed with the capability to learn specialized representations that are invariant to the influence of sensitive attributes. Consequently, our approach facilitates the reduction of biased predictions based on these attributes, fostering fairness and equitable predictions across diverse groups.

Furthermore, our approach is bolstered by the utilization of the gradient reversal layer, which plays a pivotal role in achieving robust adversarial learning. By aligning feature distributions across biased groups, the model is encouraged to minimize discrepancies in its predictions between majority and minority groups. This mechanism ensures that the model incurs substantial penalties for mispredictions on the minority group, thereby promoting fairness and equitable outcomes. The combination of specialized representations for sensitive attributes and the robust adversarial learning mechanism contributes to the model’s ability to reduce biased predictions and achieve equitable and reliable predictive performance across diverse groups, making it a promising advancement in the field of biomedical data analysis.

### 3.3. Optimizer on imbalanced data with a minimax loss

The conventional loss function based on optimization of accuracy, serves as a widely employed approach for binary and multi-class classification tasks. However, its effectiveness is limited when dealing with imbalanced biomedical data, where the presence of skewed class distributions poses significant challenges. In such scenarios, the model is susceptible to encountering a stagnation point during training, wherein all test samples are consistently predicted as the majority category. This phenomenon yields excessively high accuracy but results in very low precision or recall, indicating poor performance in classifying the minority class. Consequently, the conventional loss function fails to offer adequate guidance in addressing the inherent label imbalance, impeding the achievement of fair and equitable predictive outcomes.

To tackle the issue of label imbalance in biomedical data analysis, recent studies have focused on optimizing the Area Under the Receiver Operating Characteristic Curve (AUC), a sensible and robust metric for imbalanced-based data classification tasks^27,28^.^29^ The distinctive property of AUC lies in its aggregation across different threshold values for binary prediction, decoupling the issues of threshold setting from the model’s predictive power. Moreover, AUC takes into account both precision and recall metrics, providing a comprehensive evaluation of the model’s performance, especially in the context of biased data. Given the superior properties of AUC for handling label imbalance, it is logical to directly optimize the adversarial module based on the AUC score in our MDANN framework, instead of relying solely on the conventional accuracy-optimized cross-entropy loss function. This optimization approach allows the adversarial module to focus on improving performance in the minority class, which is crucial for enhancing fairness in the model’s predictions. To this end, Ying et al.^29^ proposed a loss function for the AUC maximization problem, we call it **minimax loss**, incorporating various parameters to facilitate the computation of the AUC score.

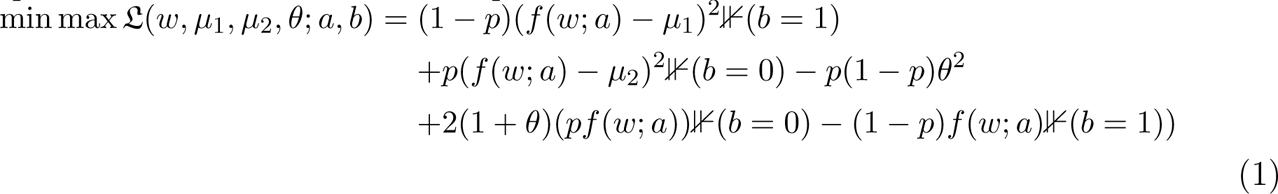

This minimax loss function incorporates various parameters, including the weights of the model *w*, the features of the samples *a*, and the labels *b*. The rate of positive samples to all samples is represented by *p*. The model’s prediction for a given sample with features *a* and weights *w* is given by *f* (*w*; *a*). Additional trainable parameters used to compute the AUC score are *µ*_1_, *µ*_2_, and *θ*, which are optimized along with the weights of the model. The loss function includes terms that contribute to the loss when the true label *b* is 1 or 0, with squared error terms measuring the difference between the model’s prediction and the parameters *µ*_1_ and *µ*_2_. These terms are weighted by (1 *− p*) and *p*, the proportions of negative and positive samples, respectively. A regularization term *−p*(1 *− p*)*θ*^2^ penalizes large values of *θ*, and additional terms adjust the model’s predictions based on *θ* and *p*. Indicator functions ⊮(*b* = 1) and ⊮(*b* = 0) ensure that each term in the loss function is only active for samples of the corresponding class. The goal of this function is to find the model weights and parameters that minimize the maximum loss over all samples, thereby improving the model’s performance on the minority class and enhancing fairness in the model’s predictions. Note that a mini-batch ADAM Stochastic Gradient Descent (SGD) method is used to update parameters for the entire MDANN. The SGD optimizer can significantly reduce the time to compute gradients at each state as well as has low chances to overfit. Additionally, ADAM has the advantage that its learning rate is adaptively adjusted, which helps all adversarial components to learn a different learning rate based on its own label distribution, especially in cases where they are responsible for learning highly biased attributes.

To efficiently update model parameters, the mini-batch ADAM Stochastic Gradient Descent (SGD) method^30^ is employed for the MDANN. ADAM’s adaptive learning rate property proves advantageous in the context of MDANN, as it enables each adversarial component to adapt its learning rate based on its specific label distribution. This adaptability is particularly valuable when adversarial components are responsible for learning highly biased attributes. By focusing on the performance of the minor class, this AUC-induced optimization methodology promotes more equitable predictive outcomes, advancing the reliability and ethical considerations of predictive models in critical medical applications.

## 4. Experimental Setup

All experiments are conducted on a cluster server with four NVIDIA RTX A5000 GPUs. We evaluate the performance of the MDANN with a varying number of adversarial components. The grid-search method is performed to fine-tune the parameters in experiments. The result under each set of experiments was averaged across five runs.

### 4.1. Datasets

In this work, we focus on two biomedical datasets. The first is from Autism Brain Imaging Data Exchange II^*^ (ABIDE II),^31^ which is a repository aggregating and openly sharing resting-state functional magnetic resonance imaging (R-fMRI) data sets with corresponding structural MRI and phenotypic information with autism spectrum disorder (ASD) and typical controls (TC). We have collected MRI imaging as well as phenotypic data from 212 individuals, with 110 ASDs and 109 TCs. The second dataset is collected from Alzheimer’s Disease Neuroimaging Initiative^**^ (ADNI) database. The ADNI was launched in 2003 as a public-private partnership, led by Principal Investigator Michael W. Weiner, MD. The primary goal of ADNI has been to test whether serial magnetic resonance imaging (MRI), positron emission tomography (PET), other biological markers, and clinical and neuropsychological assessment can be combined to measure the progression of mild cognitive impairment (MCI) and early Alzheimer’s disease (AD). We have collected 670 MRI imaging data with demographic information labeled as three categories: 350 normal cognitive (NC) samples, 152 MCI samples, and 168 AD samples. The labeling strategy employed in this study is designed to facilitate binary classification. Specifically, for the autism dataset, ASDs are labeled as 1, and TCs are labeled as 0. This labeling reflects the objective to distinguish between individuals with ASD and typical controls. In the case of the ADNI dataset, NC is labeled as 0, and both MCI and AD are collectively labeled as 1. This approach enables the differentiation between normal cognitive function and conditions indicative of cognitive impairment, including both mild cognitive impairment and Alzheimer’s disease.

In the context of demographic data, majority labels refer to specific categories or groups that may have advantages or are favored in a particular context. These labels often correspond to characteristics or attributes that are considered “normative” or “majority” within a given society or dataset. Therefore, the majority and minority groups for the two datasets are defined as shown in **Fig. 3**. For the Autism dataset, right-handedness and male are considered majority groups, mitigated by one MDANN model with two adversarial components. For the ADNI dataset, we consider aged over 78 years, educated over 18 years, and right-handedness as majority groups. Therefore, three adversarial components are needed to mitigate biases within these three attributes.

### 4.2. Evaluation metrics

To quantitatively evaluate fairness, we applied two different metrics to analyze and address fairness that would be present in the training data. We use AUC for binary classification problems, which measures the ability to distinguish between false positive rates and false negative rates. It’s important to note, however, that while AUC is a valuable measure of a model’s performance, it does not inherently provide a measure of fairness. To measure fairness, it is crucial for adversarial components to distinguish between the minority (represented as 1) and majority (represented as 0) attributes, which makes AUC a necessary metric for MDANN. We also employ the Disparate Impact (DI), which is a metric to evaluate fairness.^32^ It compares the proportion of individuals that receive a positive output for two groups: a minority and a majority group. The calculation of DI is as follows,

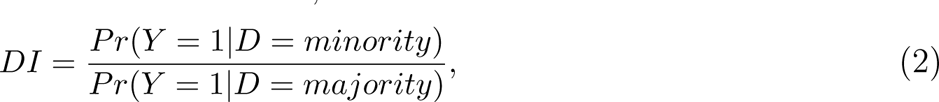

where *Pr*(*Y* = 1*|D* = *minority*) is the proportion of the minority group that received the positive outcome and *Pr*(*Y* = 1*|D* = *majority*) is the proportion of the majority group that received the positive outcome. The DI value ranges from 0 to infinity, with 1 indicating no disparate impact, meaning that the model treats all groups equally in terms of favorable outcomes. Values greater than 1 indicate a positive disparate impact, suggesting that the majority group receives more favorable outcomes than the minority group, which may indicate bias in favor of the majority group. Conversely, values less than 1 indicate a negative disparate impact, suggesting that the majority group receives fewer favorable outcomes than the minority group, which may indicate a bias against the majority group.

## 5. Results

### 5.1. Fairness enhancement achieved by the feature extractor

The input values have been fed to a pre-trained convolutional autoencoder (CAE), which is used to extract deep representations from the embedding layer. By learning to reconstruct the input, the autoencoder encourages the embedding to capture the most important and informative aspects of the data, while minimizing the impact of biased information. In MDANN, the CAE module contains several convolutional layers as the image encoder and the same number of deconvolutions layers as the decoder. Each layer employs a 5*×*5 kernel with a step size of 2 for its calculations. **Table 1** shows the prediction result of MDANN using the pre-trained CAE with different sizes of encoders and decoders. We have tested pre-trained three CAEs as the extractor of deep representations for original images. The size of deep representations is equal to the output size of the last convolutional layer in the encoder. To assess the image quality generated by the CAE modules, we adopted the Fŕechet inception distance (FID) as a metric.^33^ A lower FID score indicates an improved distribution of generated images. Experiments show that CAE-2 could reach the maximum accuracy of 0.73 and averaged DI of 0.77 for sensitive attributes for AD prediction. In contrast, MDANN achieves only 0.63 accuracy and 0.68 averaged DI when utilizing original data without any feature extraction (as depicted in the first row of **Table 1**). This substantiates the assertion that deep representations encapsulate an enriched trove of information within higher-dimensional spaces, thereby enhancing the model’s fairness in prediction.

**Table 1.** Performance of MDANN using different CAE modules.

| Module | Embedding Size | # of Conv layers | FID score | Accuracy | Averaged DI |
| --- | --- | --- | --- | --- | --- |
| N/A | N/A | N/A | N/A | 0.63 | 0.68 |
| CAE-1 | $64 \times 64 \times 128$ | 1 | 556.74 | 0.63 | 0.67 |
| **CAE-2** | **$32 \times 32 \times 256$** | **2** | **490.98** | **0.73** | **0.77** |
| CAE-3 | $16 \times 16 \times 512$ | 3 | 555.54 | 0.66 | 0.72 |

Furthermore, more embedding layers might enable better reconstruction, preserving more details of the input data. However, this improvement does not always translate into better performance for specific downstream tasks. When the number of convolutional layers is increased to 3 (CAE-3), we got a lower FID score than the score of CAE-2. Thus, the CAE-2 has been selected as the best feature extractor for the following experiments. Illustrated in **Fig. 2** are the generative images of CAE-2 during a 100-epoch training process for a single sample image. Notably, the output image converges satisfactorily over the course of these epochs. In order to obtain a stable deep representation, CAE training was extended to 200 epochs.

**Fig. 2.**
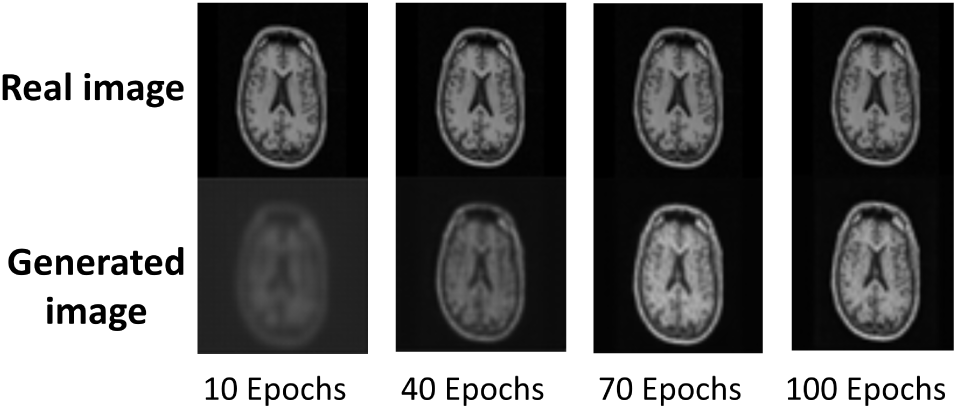
The training results of CAE-2 module. As illustrated by the generated images for one real sample image with 10, 40, 70 and 100 epochs respectively. The trained model is prone to converge at epoch 100.

### 5.2. Improved optimization strategies to handle label imbalance

Since each sensitive label for training is imbalanced, (i.e., 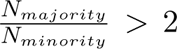) we can further explore if all ADs collaboratively optimize the loss function to learn the best model parameter against label imbalance. **Fig. 3** shows the specific label distribution of each sensitive feature in two datasets. It can be obverse that the number of majority groups is greater than that of minority groups. Generally, the label-imbalance problem could lead to a biased prediction. Specifically, the machine learning models are prone to predict the minority class as the majority class. Thus, we have tested and plotted the performance of two prediction tasks using the same model with two different loss functions (minimax loss and standard cross entropy loss function as a comparison) during the training process, as shown in **Fig. 4**. Results of the prediction utilizing different losses and optimizers are shown as two rows, ACC-induced Cross-Entropy Loss and AUC-induced Min-max Loss. Each row contains four columns, showing the classification performance (ACC) for each sensitive feature and the global AUC score for the predictor. AUC results show that a minimax loss does improve the classification performance while using cross-entropy loss could only achieve around 0.6 AUC score, tending to be less distinguishable between minority (represented as 1) and majority (represented as 0) attributes. Particularly, the prediction accuracy of two categories for handedness converges at 0.6 since it did not learn any relation between features and labels due to the high-imbalanced data (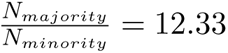). The second row shows that the weighted min-max loss overcame the label-imbalanced problem, where the AUC score increased to 0.8 demonstrating that employing a minimax loss significantly increases the accuracy for minor classes. The predictor gained an accuracy of 0.8 for patients who were left-handed while was only 0.6 if using a classical cross-entropy loss. Note that hyperparameters and model architecture are kept the same except for the loss function. These results support that we have implemented an effective approach to address the label-imbalanced issue which is a big concern for biomedical data.

**Fig. 3.**
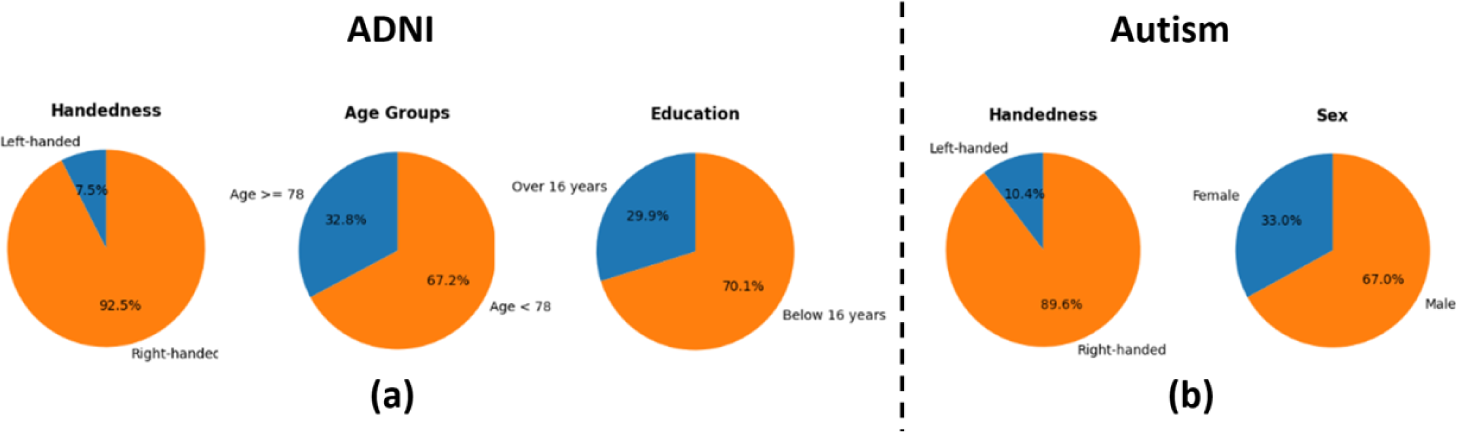
Demographic Distribution in Autism and ADNI datasets. (a). the distribution of Handedness and Sex in the Autism dataset. The left chart illustrates the proportion of left-handed and right-handed individuals, while the right chart delineates the gender distribution. (b). the distribution of Age Groups, Handedness, and Education in the ADNI dataset. The left chart categorizes individuals into two age groups, the middle chart shows the distribution of left-handed and right-handed individuals, and the right chart divides the population into two education levels. The majority groups are represented in orange color and minority groups are in blue color.

**Fig. 4.**
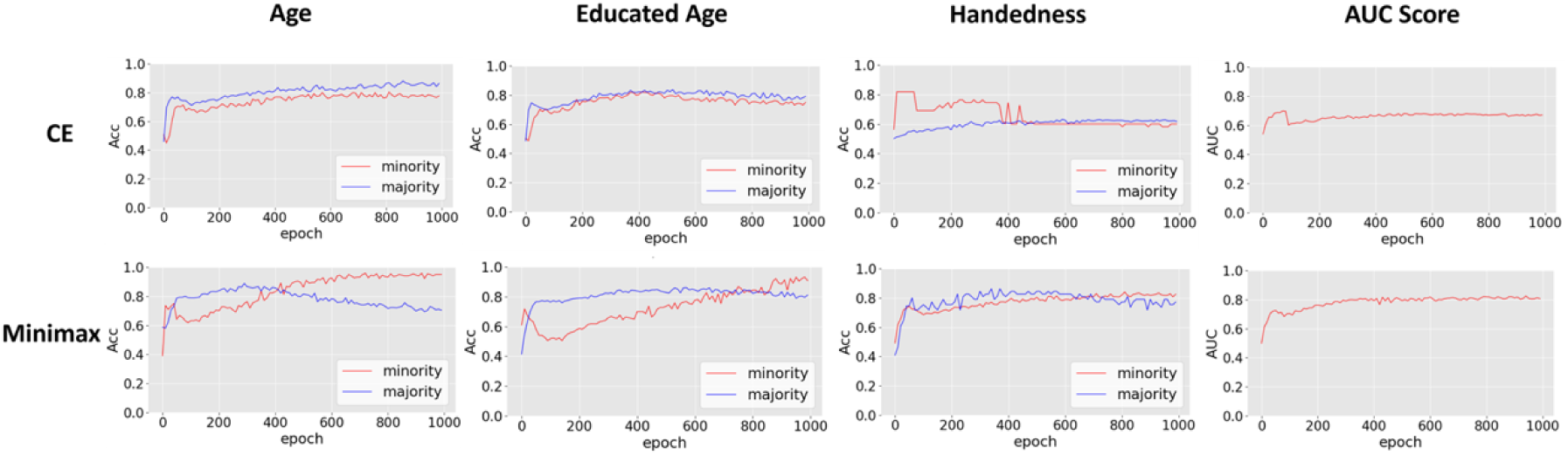
Prediction results addressing the label-imbalance problem. Three sensitive attributes from ADNI datasets were included: age (with aged over 78 years as the minority group), educated years (with educated over 16 years as the minority group), and handedness (with left-handed as the minority group). Rows represent the prediction metrics with different loss functions: cross-entropy loss and minimax loss. The first three columns show the accuracy of the predictor when predicting sensitive features respectively, and the last column shows the AUC score of the predictor.

### 5.3. Bias mitigation via multiple components in the adversarial module

Next, we investigated the impact of the number of adversarial components on prediction performance in the context of fairness enhancement. To do so, we trained the MDANN while simultaneously addressing multiple sensitive features for AD prediction tasks. We systematically compared three variants of the MDANN with differing numbers of adversarial modules. **Fig. 5** depicts the predictive performance. In this figure, panel (a) illustrates the accuracy and AUC score of predictors employing different numbers of adversarial components while incorporating cross-entropy loss into the MDANN. The red line represents the baseline, which corresponds to regular training without an adversarial module. When optimizing the model using cross-entropy loss, we observed that increasing the number of adversarial components led to a reduction in prediction performance, as evidenced by diminished accuracy and AUC scores. Specifically, the lowest accuracy of 0.65 was obtained for disease prediction when the MDANN simultaneously mitigated four attributes. This outcome is attributed to the inclusion of more imbalanced features during the training process, which the cross-entropy loss cannot effectively address. Additionally, **Fig. 5** (b) illustrates the accuracy and AUC score when employing minimax loss for training. We observed that during regular training (baseline), the accuracy initially reached a peak and then declined, eventually stabilizing at an accuracy of 0.76 over 1000 epochs. This behavior reflects the outcome of the AUC-induced minimax loss, given its objective of maximizing the AUC, and thereby precluding a guarantee that accuracy is consistently optimized. Fortunately, increasing the number of adversarial components proved effective in mitigating this issue. As the number of imbalanced labels increased, the number of negative gradients propagated backward also increased, leading to a reduction in the convergence speed. Consequently, employing more adversarial components overcame this challenge and effectively combined the advantages of the loss function, ultimately achieving the highest accuracy of 0.85 for AD prediction.

**Fig. 5.**
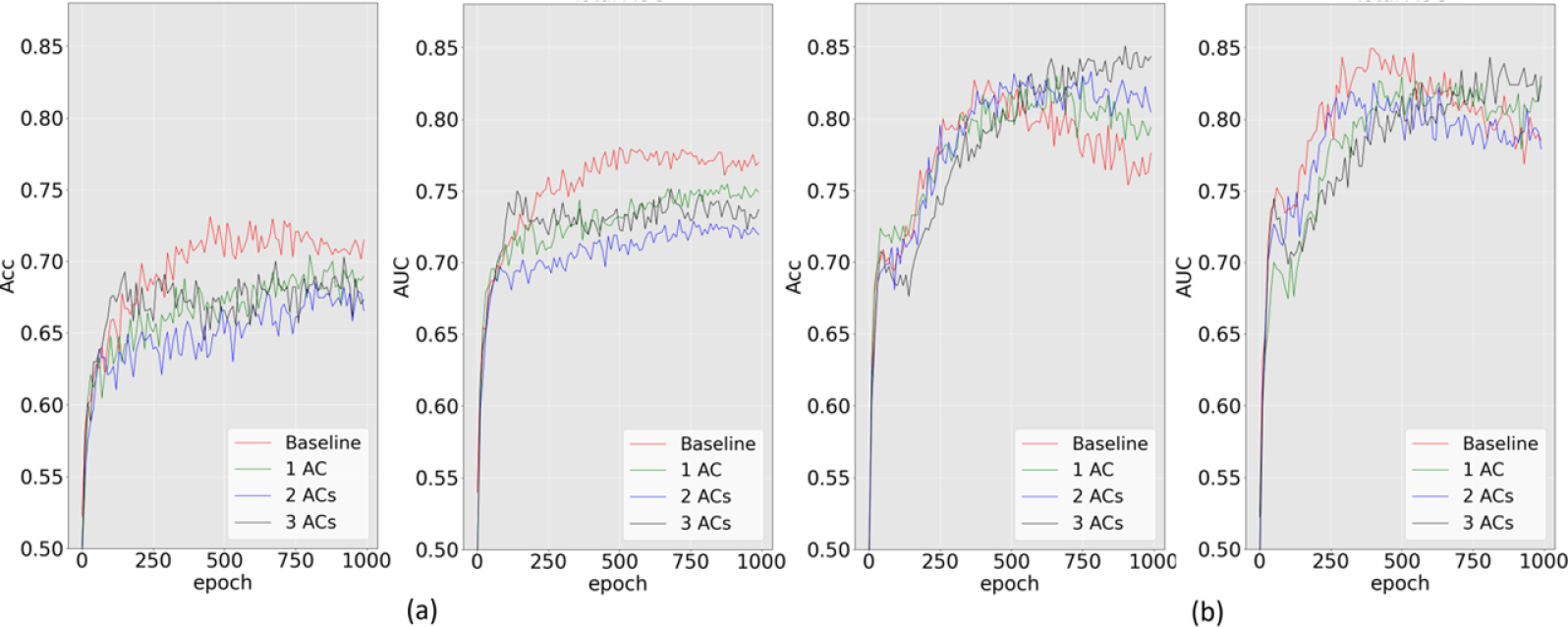
Validation accuracy (Acc) and AUC score for ADNI disease prediction under two scenarios. (a). MDANN trained with cross-entropy loss. (b). MDANN trained with minimax loss. Three biased attributes: age, handedness, and educated years, were used as three embedded adversarial components (ACs) for bias mitigation. The baseline was the regular machine learning model trained without any AC. All images have been fed to a pre-trained CAE with two convolutional layers (with deep representations size of 32*×*32*×*256).

### 5.4. Combined mechanisms for fairness enhancement

Having elucidated the individual contributions of the feature extractor, minimax loss, and adversarial module through experimental data, we now turn our attention to a comprehensive evaluation of these components within the MDANN framework. We analyzed three fair-enhancing mechanisms applied to biomedical datasets, and we present their efficacy as modules in the MDANN. To investigate MDANN’s potential in enhancing fairness, we performed experiments to combine these modules in an integrated manner. We tested the MDANN while simultaneously addressing multiple sensitive features for Autism prediction tasks. A comparison was conducted across various approaches, including two variants of MDANN with differing numbers of adversarial components, and two other methods referred to as Zhang et al.^34^ and Ganin et al.^22^ The examination encompassed two sensitive features: sex and handedness.

**Fig. 6** depicts the performance of different models in alleviating biases and enhancing fairness in predictions. For the sensitive feature of sex, Zhang et al. yielded an accuracy of 0.479, an AUC of 0.5, and a DI of 0.386. These results indicate both poor predictive performance and significant bias, suggesting limitations in the method’s ability to generalize across multiple domains and mitigate bias effectively to enhance fairness for biomedical data. Ganin et al. showed improvement with an accuracy of 0.63, an AUC of 0.63, and a DI of 0.673, but still exhibited shortcomings in handling disparities between the majority groups and minority groups. While the proposed MDANN method demonstrated superior predictive performance and fairness with an accuracy of 0.71, an AUC of 0.73, and a DI of 0.841 for the variation with 2 adversarial components. The results reflect the MDANN’s ability to differentiate between classes and correctly classify instances, as well as its commendable increase in fairness.

**Fig. 6.**
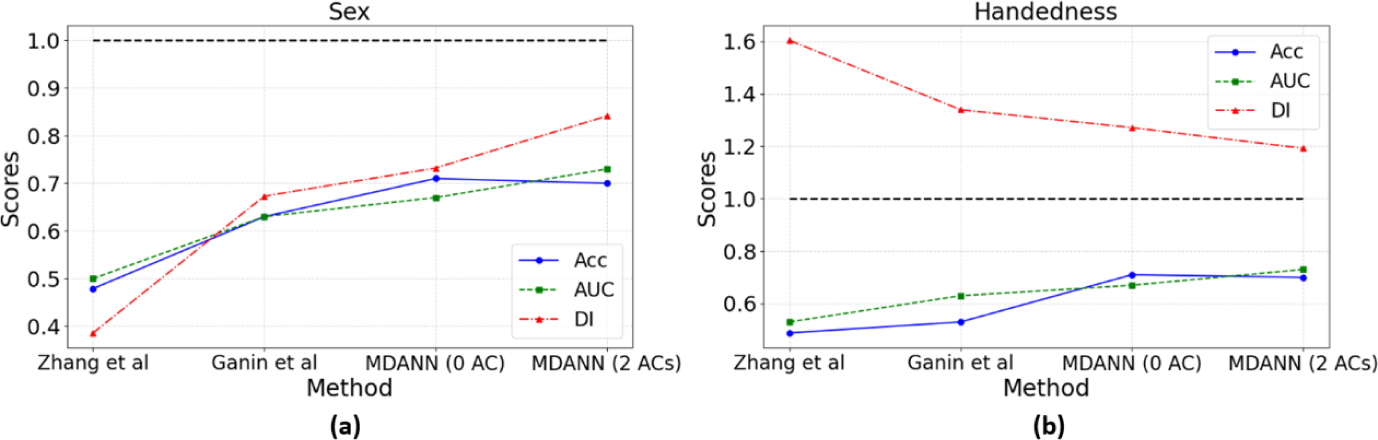
Comparative performance of different methods on sex and handedness. Four methods have been tested: Zhang et al., Ganin et al., MDANN with 0 adversarial components (AC), and MDANN with 2 ACs, across two sensitive features: (a) Sex and (b) Handedness. The three performance metrics are represented by different line styles and markers: Accuracy (ACC) is shown in blue with circular markers and a solid line, Area Under the Receiver Operating Characteristic Curve (AUC) is depicted in green with square markers and a dashed line, and Disparate Impact (DI) is illustrated in red with triangular markers and a dash-dot line. A horizontal black dashed line at y=1 serves as a reference for the DI metric, where values closer to 1 indicate enhanced fairness. The results highlight the superior performance of MDANN, particularly the variation with 2 ACs, in terms of both predictive accuracy and fairness.

A similar pattern emerged for the mitigation of handedness. Zhang et al. performed the worst, with an accuracy of 0.488, an AUC of 0.53, and a DI of 1.604, reflecting the method’s inability to adequately address the challenges of fairness enhancement. Ganin et al. improved on these figures but were still surpassed by MDANN, which achieved an accuracy of 0.70, an AUC of 0.73, and a DI of 1.192 for the variation with 2 adversarial components.

## 6. Conclusion

In this study, we introduced an MDANN framework, which incorporates three modules to perform fair prediction on biomedical imaging datasets. Our empirical findings highlight the efficacy of adversarial modules in the MDANN framework, effectively mitigating biases and promoting fairness by addressing multiple sensitive features. Additionally, the utilization of AUC-based minimax loss functions demonstrates their superior handling of label imbalance, leading to improved classification performance for the minority class. Furthermore, we showcase the potential of deep representations extracted from a pre-trained convolutional autoencoder, resulting in enhanced prediction accuracy and fairness in AD and Autism prediction. The investigations underscore the promising potential of MDANN as a method for fairness enhancement, particularly in scenarios where the sensitive attributes of sex and handedness are of concern, simultaneously. Our experimental results provide intuition into the reasons for MDANN’s superiority, reflecting its potential as a valuable method for enhancing both predictive performance and fairness. Nonetheless, there are future opportunities for extending and enhancing MADNN. First, we will use a generative adversarial network (GAN)^35^ for feature extraction rather than CAEs. Second, it should be recognized that the current structure of adversarial components is simple and uniform in MDANN. We will extend MDANN by integrating adversarial components with distinct architectures. Such an approach would enable the training of specialized components tailored to each sensitive feature, accommodating its unique distribution characteristics.

## Acknowledgments

Preprint of an article submitted for consideration in Pacific Symposium on Biocomputing © [2024] World Scientific Publishing Co., Singapore, http://psb.stanford.edu/. We would like to extend our gratitude to the Autism Brain Imaging Data Exchange (ABIDE). We appreciate the contribution of Chen Song, who provided the Alzheimer’s Disease Neuroimaging Initiative (ADNI) dataset used in our experiments. Data collection and sharing for this project was funded by the ADNI (National Institutes of Health Grant U01 AG024904) and DOD ADNI (Department of Defense award number W81XWH-12-2-0012). ADNI is funded by the National Institute on Aging, the National Institute of Biomedical Imaging and Bioengineering, and through generous contributions from the following: AbbVie, Alzheimer’s Association; Alzheimer’s Drug Discovery Foundation; Araclon Biotech; BioClinica, Inc.; Biogen; Bristol-Myers Squibb Company; CereSpir, Inc.; Cogstate; Eisai Inc.; Elan Pharmaceuticals, Inc.; Eli Lilly and Company; EuroImmun; F. Hoffmann-La Roche Ltd and its affiliated company Genentech, Inc.; Fujirebio; GE Healthcare; IXICO Ltd.; Janssen Alzheimer Immunotherapy Research & Development, LLC.; Johnson & Johnson Pharmaceutical Research & Development LLC.; Lumosity; Lundbeck; Merck & Co., Inc.; Meso Scale Diagnostics, LLC.; NeuroRx Research; Neurotrack Technologies; Novartis Pharmaceuticals Corporation; Pfizer Inc.; Piramal Imaging; Servier; Takeda Pharmaceutical Company; and Transition Therapeutics. The Canadian Institutes of Health Research is providing funds to support ADNI clinical sites in Canada. Private sector contributions are facilitated by the Foundation for the National Institutes of Health (www.fnih.org). The grantee organization is the Northern California Institute for Research and Education, and the study is coordinated by the Alzheimer’s Therapeutic Research Institute at the University of Southern California. ADNI data are disseminated by the Laboratory for Neuro Imaging at the University of Southern California. XJ is CPRIT Scholar in Cancer Research (RR180012), and he was supported in part by the Christopher Sarofim Family Professorship, UT Stars award, UTHealth startup, the National Institute of Health (NIH) under award number R01AG066749, R01AG066749-03S1, R01LM013712, U24LM013755 and U54HG012510, and the National Science Foundation (NSF) #2124789.

## Footnotes

* ABIDE II data used in paper are available on http://fcon1000.projects.nitrc.org/indi/abide/abideII.html

** Data used in preparation of this article were obtained from the Alzheimer’s Disease Neuroimaging Initiative (ADNI) database (adni.loni.usc.edu). As such, the investigators within the ADNI contributed to the design and implementation of ADNI and/or provided data but did not participate in analysis or writing of this report. A complete listing of ADNI investigators can be found at: http://adni.loni.usc.edu/wp-content/uploads/howtoapply/ADNIAcknowledgementList.pdf

## Notes

### Competing Interest Statement

The authors have declared no competing interest.

### Summary of Updates

Add reference and acknowledgment of ANDI data usage.

